# Identification of midbrain neurons essential for vocal communication

**DOI:** 10.1101/310250

**Authors:** Katherine Tschida, Valerie Michael, Bao-Xia Han, Shengli Zhao, Katsuyasu Sakurai, Richard Mooney, Fan Wang

## Abstract

Vocalizations are an essential medium for communication and courtship in numerous mammalian species ranging from mice to humans. In mammals, the midbrain PAG serves as an obligatory node in a vocalization-related network that spans the forebrain and brainstem^1–3^, as bilateral lesions of the PAG result in mutism^2–5^. Despite the PAG’s importance for vocal production, the identity, function, and connectivity of PAG neurons involved in vocalization has remained elusive, in part because the PAG is a functionally and anatomically heterogeneous structure that serves myriad roles including nociception, defensive behaviors, and autonomic regulation^6–9^. Here we used a viral genetic “tagging” method^10,11^ to identify a distinct subset of PAG neurons in the male mouse that are selectively activated during the production of ultrasonic vocalizations (USVs) elicited by female cues. Silencing these PAG-USV neurons rendered males mute without affecting their other courtship behaviors and also impaired their ability to attract female mice in a social choice assay. Activating these neurons using chemogenetic or optogenetic methods strongly elevated USV production, even in the absence of female cues. Notably, the timing of individual USVs was entrained to the expiratory phase of breathing but not to the pattern of optogenetic stimulation, suggesting that PAG-USV neural activity initiates and sets the duration of vocal bouts and recruits downstream premotor circuits that precisely pattern vocal output. Consistent with this idea, we found that PAG-USV neurons extend axons into pontine and medullary regions that are speculated to contain premotor central pattern generators important for vocalization^3,12,13^. These experiments establish the identity of the PAG neurons selectively required for USV production in mice, map their efferent connections, and demonstrate the communicative salience of male USVs in promoting female social affiliation.

## Results

The functional heterogeneity of the PAG has precluded the identification of the exact subset of PAG neurons important for vocal control. Therefore, a critical step to dissect the brain-wide networks important for vocalization requires a method to selectively identify and manipulate vocalization-related PAG neurons. To this end, we used a recently developed immediate early gene Fos-based method (CANE^10,11^; Fig. 1) to genetically label, identify, and manipulate PAG neurons that are selectively active when male mice produce courtship ultrasonic vocalizations (USVs), which comprise individual vocal elements (i.e., syllables) that are organized into multisyllabic bouts that last from hundreds of milliseconds to tens of seconds in duration^14–19^ (Fig. 1A). As a prelude to using CANE, we established that Fos expression increased in a subset of neurons in the caudolateral PAG in single-housed male mice that produced USVs to a female social partner or to female urine, but remained at baseline levels in males that failed to vocalize to female cues or in the presence of control odors that did not elicit vocalization (Fig. 1B-C). Across all mice, the number of Fos-positive neurons in the caudolateral PAG positively correlated with the number of USVs produced (Fig. 1D), tentatively identifying PAG neurons important to USV production (i.e., PAG-USV neurons).

**Figure 1.**
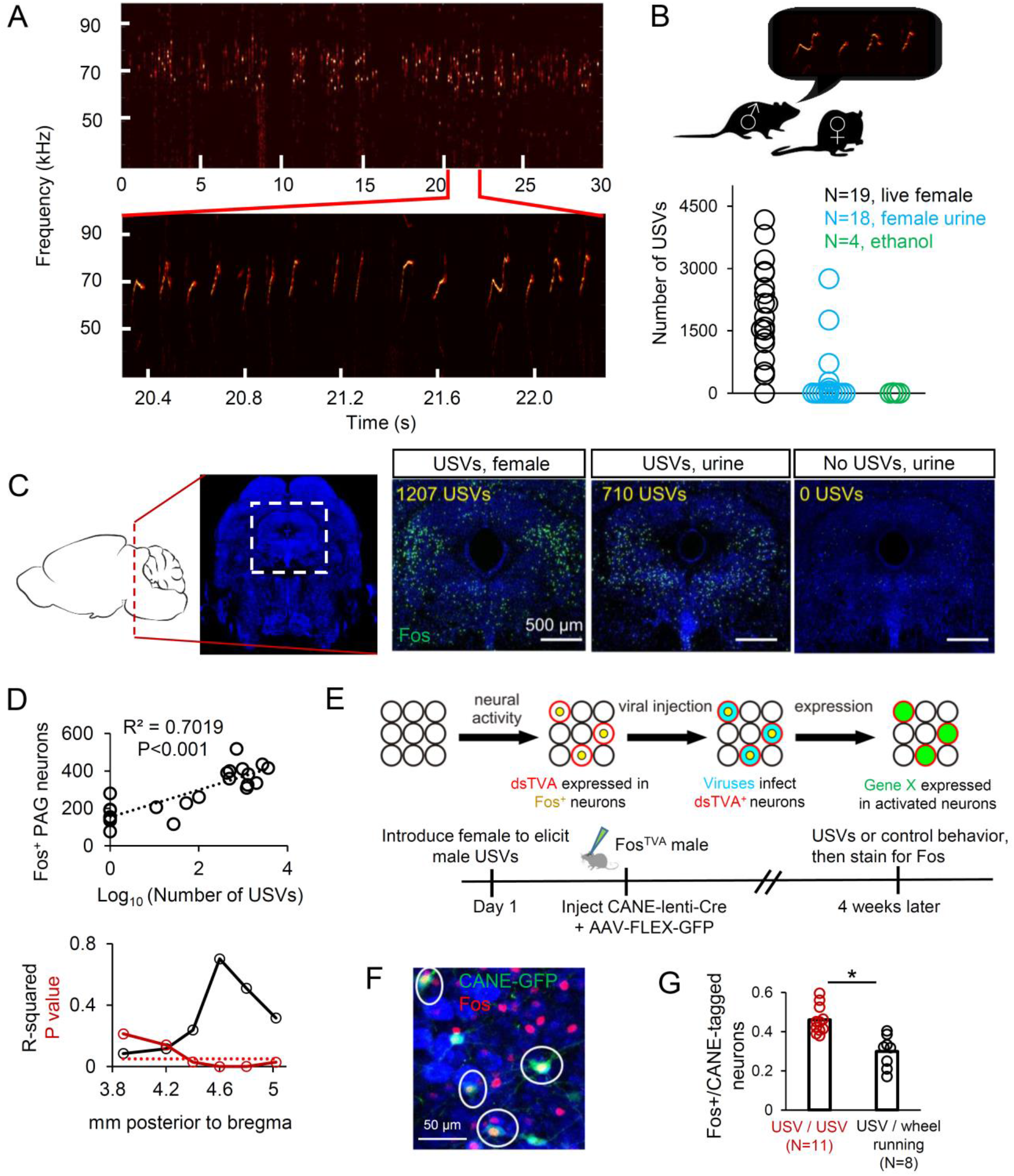
The caudolateral PAG contains a subset of neurons that are active during USV production. (A) Spectrogram of USVs produced by a male mouse during a social encounter with a female. Inset shows expanded view. (B) (Top) Schematic showing male mouse producing courtship USVs toward a female social partner. (Bottom) Quantification of number of USVs produced by male mice during encounters with female social partners (black), female urine (blue), or a non-social odor, ethanol (green). (C) Left-most panels show location of the caudolateral PAG in sagittal and coronal section. Three right-most panels show representative confocal images of Fos expression in the caudolateral PAG following social experience with a female social partner accompanied by USV production, female urine accompanied by USV production, and female urine without USV production (green, Fos; blue, NeuroTrace). (D) (Top) The total number of Fos-positive neurons in the caudolateral PAG is positively correlated with the number of USVs produced by different males. Quantification is shown for the same plane of section as in (C). (Bottom) The strength (R^2^) and significance (p value) of the linear regression between the number of USVs produced (log10) and Fos expression in the caudolateral PAG is plotted across a 1.2 mm rostocaudal extent of the PAG (N=10 males given female social partner; N=10 female urine; N=2 ethanol). (E) Schematic of the CANE method (top) and the experimental time line to permanently express transgenes in PAG-USV neurons using CANE (bottom). (F) Confocal image showing overlap between CANE-GFP-labeled PAG-USV neurons (green) after a USV production toward a female and Fos (red) induced by a subsequent vocal encounter with a female. (G) Quantification of the overlap between CANE-GFP-labeled PAG-USV neurons and Fos expression elicited by USVs (red points, N=11 mice) or by wheel running in the home cage (black, N=8 mice; p<0.001 for difference between groups, Mann Whitney U test).

We used CANE to express GFP in PAG-USV neurons, allowing us to map their axonal projections and also establish whether they represent a stable neuronal population. Briefly, we elicited USVs from single-housed male Fos^TVA^ mice (during a 30-60 min session with a female), then injected the caudolateral PAG with a combination of CANE-lenti-Cre and AAV-FLEX-GFP viruses (Fig. 1E). This approach extensively labeled PAG-USV neurons with GFP, revealing that their axons terminate in a variety of targets in the midbrain, pons, and medulla (described in more detail below). We also established that ~50% of PAG neurons labeled with GFP following an encounter in which the male emitted USVs toward a female were re-activated (i.e., upregulated Fos) following a second round of vocalization ~4 weeks later (Fig. 1F-G, 46.0 ± 2.1%, N=11 mice). This proportion of reactivated cells is significantly greater than the number of double-labeled PAG neurons observed when the first vocal social encounter was followed 4 weeks later by a 30-60 min period of wheel-running in the mouse’s home cage (Fig. 1G, 29.9 ± 2.9%, N=8, p<0.001, Mann Whitney U test). Therefore, many of the same PAG-USV neurons are active during vocal bouts separated by many weeks, indicating that a stable and distinct subpopulation of PAG neurons is activated during USV production.

To test the causal role of PAG-USV neurons in USV production, we used CANE to express tetanus toxin light chain (TeLC) in PAG-USV neurons, which over days to weeks irreversibly blocks their ability to release neurotransmitters (see Methods; CANE-lenti-Cre and AAV-FLEX-TeLC injected bilaterally into the caudolateral PAG of a Fos^TVA^ male that had recently vocalized). Blocking transmitter release from PAG-USV neurons strongly suppressed male USV production to nearby females: most (7/11) males were rendered completely mute for USV production, while the remaining 4 males emitted extremely low levels of USVs, and the extent of this vocal suppression was well related to the number of TeLC-expressing PAG-USV neurons (Fig. 2A-B; 2 weeks after injection, mean USV counts were 4.7 ± 2.3 % of pre-injection level; Movies S1-2).

**Figure 2.**
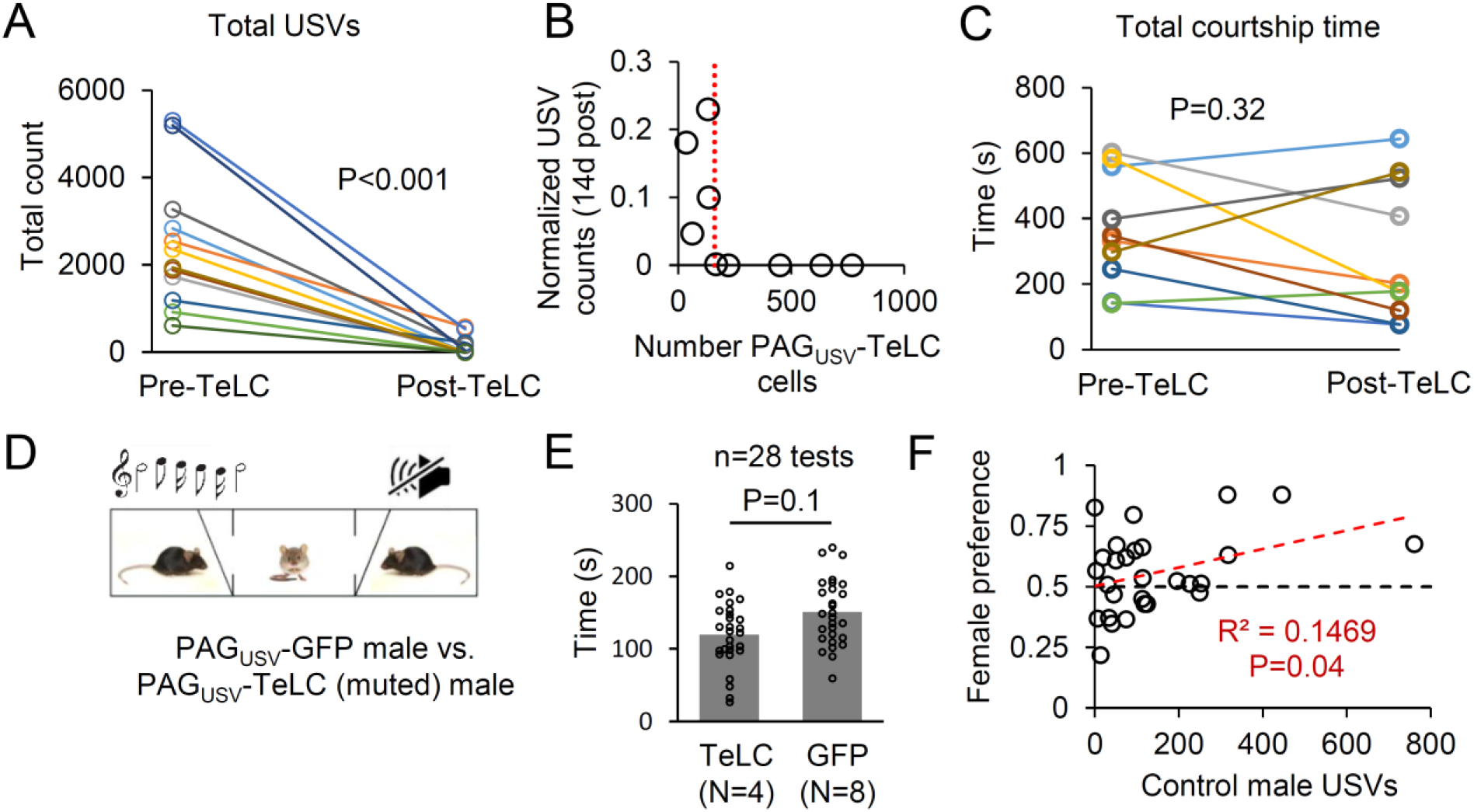
The production of courtship USVs requires PAG-USV neurons and promotes female social affiliation. (A) Blocking neurotransmitter release from PAG-USV neurons with CANE-driven expression of TeLC abolishes the production of male courtship USVs (N=12 mice, p<0.001, Wilcoxon signed-rank test). (B) The number of TeLC-expressing PAG-USV neurons is plotted against the normalized USV count for each PAG_USV_-TeLC male (total day 14 USVs/total day 1 USVs; post-hoc histology quantified for N=9 mice). (C) Blocking neurotransmitter release from PAG-USV neurons has no significant effect on total time spent courting a female (p=0.32, Wilcoxon signed-rank test). (D) Schematic showing the set-up for the three-chambered test, comparing female preference for a muted PAGusv-TeLC male versus a freely vocalizing PAG_USV_-GFP male. (E) Females tend to spend more time with control, vocalizing males than with muted males (p=0.1, Wilcoxon signed-rank test, N=28 tests from N=4 PAG_USV_-TeLC mice each paired against 1-3 PAG_USV_-GFP controls (N=8 in total), with each TeLC/GFP pair tested with 2-4 females). (F) Vocal output of the control PAG_USV_-GFP male is plotted against female preference for the control male (>0.5 is preference for control male)). R^2^ and p value shown for linear regression.

Although these findings are consistent with the idea that PAG-USV neurons are directly involved in USV production, silencing PAG-USV neurons could indirectly block vocalization, perhaps by decreasing social or sexual motivation. To control for this possibility, we compared the amount of time PAG_USV_-TeLC males spent courting a female before and 2 weeks after viral injection (Fig. 2C). We noted that the amount of time spent engaged in courtship tended to decrease on average post-injection (83.1 ± 16.5% pre-injection level, see Ext. Data Fig. 1 for low vs. high intensity courtship). However, this decrease was not significant (Fig. 2C, Ext. Data Fig. 1) and was not different from the change seen over 2 weeks in PAG_USV_-GFP control males (81.8 ± 6.3% pre-injection courtship level, N=14 mice). Therefore, PAG-USV neurons are necessary for USV production but are not required for normal levels of male non-vocal courtship behavior.

We next took advantage of our ability to selectively abolish USVs to shed light on whether a male’s ability to vocalize influences his ability to attract females. Because males normally produce USVs in concert with non-vocal courtship behaviors^18–21^, it has been challenging to determine whether male USVs influence female social affiliation independently of non-vocal courtship. To address this issue, we tested female preference for muted TeLC-expressing males versus control GFP-expressing males in a three-chambered preference test, during which the control male can freely vocalize, but neither male can engage in non-vocal courtship behaviors (Fig. 2D). We found that females tended to prefer control, vocal males over muted males (Fig. 2E, p=0.1, Wilcoxon signed-rank test), but when we considered the amount that each control male had vocalized, we found that the degree of female preference for the vocal male increased significantly with increasing vocal output (Fig. 2F, p=0.04 for linear regression, >0.5 is preference for control male, N=28 tests). These findings show that male USVs can bias female social preference independently of other courtship behaviors, thus establishing a role for courtship USVs in promoting female social affiliation.

The present finding that PAG-USV neurons are necessary for USV production raises the possibility that activating these neurons is sufficient to elicit USV production in the absence of any social interactions or social cues. To test this idea, we used CANE to bilaterally express the excitatory DREADDs receptor hM3Dq in PAG-USV neurons (Fig. 3A). Six weeks later, we administered either CNO or saline (i.p.) and measured the vocal behavior of individual males while alone in a novel test chamber, a condition in which males normally emit few USVs (Fig. 3A, 9.3 ± 7.4 USVs over 60 min., N=4). After CNO treatment, almost all (7/8) hM3Dq-expressing males dramatically increased their USV production (Fig. 3A; 279.4 ± 138.6 USVs over 60 min., p=0.02, Wilcoxon signed-rank test), whereas after saline treatment, they emitted USVs at low rates similar to controls and CNO-treated males that did not express the hM3Dq receptor (Fig. 3A, saline day: 17.0 ± 7.0 USVs, CNO only males: 1.5 ± 0.9 USVs over 60 min., N=4 mice). To examine the linkage between PAG-USV activation and USV production with greater temporal precision, we then used CANE to express channelrhodopsin (ChR2) in PAG-USV neurons and implanted fiber optics bilaterally over the caudolateral PAG. Optogenetically activating PAG-USV neurons with blue light was sufficient to rapidly elicit USVs in isolated males, indicating that activation of PAG-USV neurons is tightly coupled to USV production (Fig. 3B-C, trains: 5-10 Hz, 1-2s duration; tonic pulses: 0.5-2s duration; min. latency from laser onset to first USV was 23.4 ± 8.6 ms, mean latency was 406.6 ± 0.5 ms, n=104 laser stimuli from N=2 PAG_USV_-ChR2 mice). We note that the acoustic features of optogenetically-elicited USVs were similar to natural courtship USVs recorded from the same males, with subtle changes to certain acoustic parameters that fell within the normal range seen for courtship USVs, while chemogenetically-elicited USVs also resembled natural courtship USVs but had more notable acoustic differences (Fig. 3C and Ext. Data Fig. 2). These findings extend previous studies showing that non-selective activation of the PAG can elicit vocalization^2–4^ and establish that activation of a small (<500) and selective subset of PAG neurons is sufficient to elicit USV production in socially isolated male mice.

**Figure 3.**
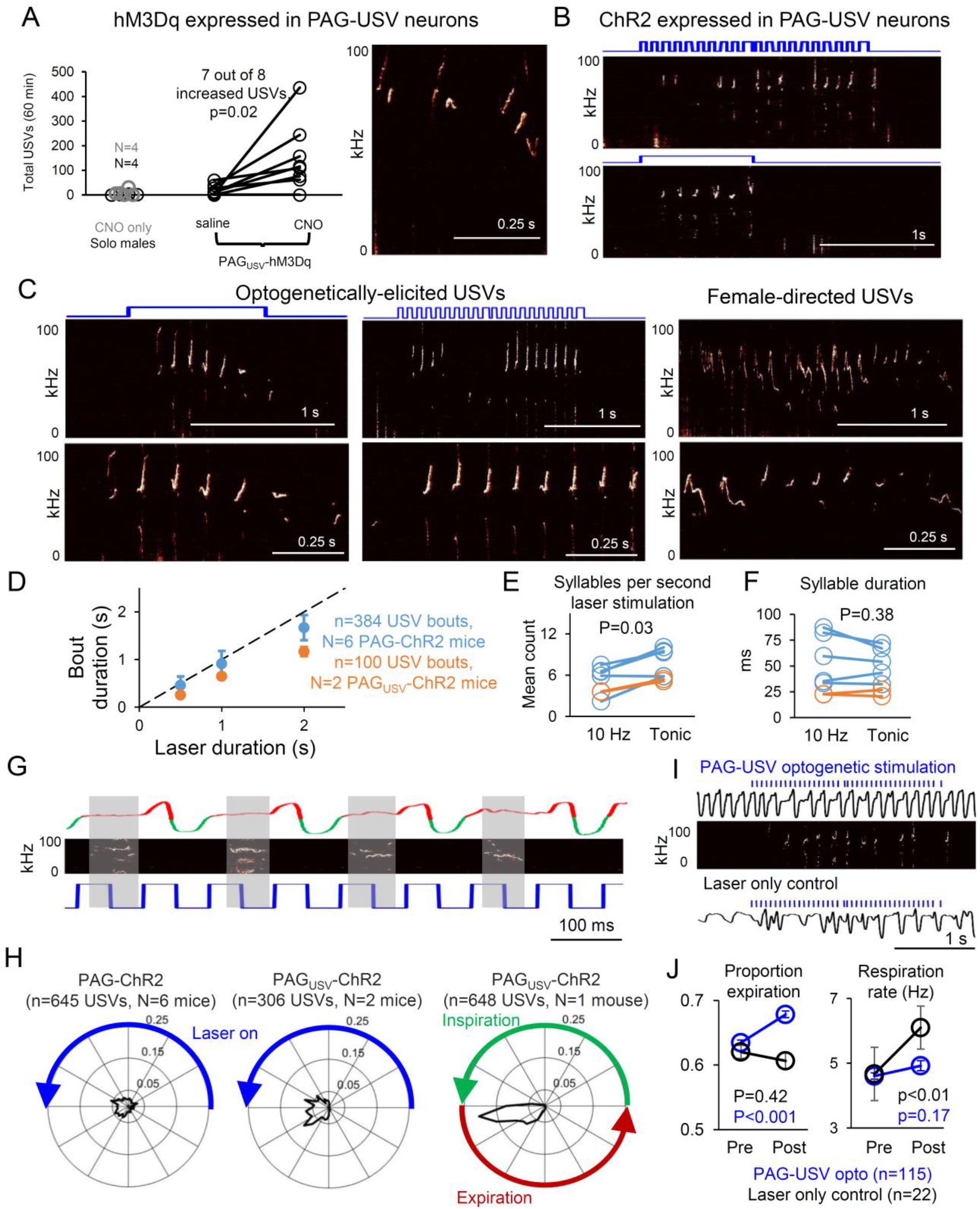
PAG-USV neuronal activity is sufficient to elicit USV production in the absence of social cues, initiates USV bouts, and determines vocal bout duration. (A) Total USVs produced in a 60 min. solo test period are shown for different groups of males in chemicogenetic experiments: control males (no virus, no CNO, gray points, N=4), males not expressing hM3Dq but treated with CNO (black, N=4), and males with CANE-driven expression of hM3Dq in PAG-USV neurons that were treated with saline (i.p., N=8) or CNO (p=0.02 for PAG_USV_-hM3Dq mice saline versus CNO USVs, Wilcoxon signed-rank test). (Right) Spectrogram showing USVs produced by a CNO-treated PAG_USV_-hM3Dq male. (B) Optogenetic activation of PAG-USV neurons is sufficient to elicit USVs from males in the absence of female cues. Note that patterned 10Hz (top) and tonic (bottom) laser stimuli both elicited multisyllabic bouts of USVs. (C) Spectrograms comparing optogenetically-elicited USVs (left and middle) and female-directed USVs (right) from the same male. Bottom panels show expanded views from the top panels. (D) The duration of optogenetically-elicited bouts of USVs is similar to the duration of laser stimuli used to activate PAG-USV neurons (n=384 opto-USV bouts from N=6 PAG-ChR2 mice; n=100 opto-USV bouts from N=2 PAG_USV_-ChR2 mice, mean ± SE). (E) Comparison of the number of USV syllables elicited per second of laser stimulation is shown for 10 Hz versus tonic laser stimuli for PAG-ChR2 mice (blue) and PAG_USV_-ChR2 mice (orange, p=0.03 for difference between 10 Hz and tonic, Wilcoxon signed-rank test). (F) Same as (E), except comparing mean syllable duration for USVs optogenetically elicited by 10 Hz and tonic laser stimuli (p=0.38). (G) A representative portion of a bout of optogenetically-elicited USVs is shown for a PAG_USV_-ChR2 mouse, with breathing shown in the top trace (inspirations are downward deflections shown in green, expiration is shown in red, see Methods), USVs shown in the spectrogram (middle), and 10 Hz laser stimulus shown in blue (bottom). Gray shading during USV times shows the clear alignment between USVs and expiration. (H) (Left, middle) Polar plots showing the distribution of onset times of individual optogenetically-elicited USV syllables relative to the duty cycle of the preceding laser pulse (10 Hz, 50ms on and 50ms off) for PAG-ChR2 mice (left) and PAG_USV_-ChR2 mice (right). Blue line indicates the laser-on portion of the laser duty cycle. Radial values (ranging from 0 to 0.25) represent proportion of total observations at a given time in the laser duty cycle, and total area inside the shaded black line is equal to one. (Right) Polar plot showing the distribution of onset times of individual optogenetically-elicited USV syllables relative to the respiratory cycle (green line is inspiration, red line is expiration). USV onsets are tightly entrained to the respiratory cycle, occurring just after the end of inspiration (n=648 USVs). (I) Representative breathing traces are shown for a PAG_USV_-ChR2 mouse during optogenetic activation of PAG-USV neurons (top, breathing shown in black, laser in blue, USVs in spectrogram) and during a control period in which the laser was turned on in the test chamber but was disconnected from the optogenetic ferrule (bottom; breathing, black; laser, blue). (J) Optogenetic activation of PAG-USV neurons caused a significant increase in the proportion of the respiratory cycle occupied by expiration (left, blue, n=115 epochs of laser stimulation, p<0.001, Wilcoxon signed-rank test) and no change in respiration rate (right, blue). When the laser was not connected to the ferrule, laser light alone did not change the expiration/inspiration ratio (left, black, n=22 epochs of laser stimulation, p=0.42) and caused a significant increase in breathing rate (right, black, p<0.01), likely due to a startle response (mean ± SE are plotted for all panels of J).

Electrical or chemical stimulation of the PAG elicits vocalization in a variety of vertebrates^2–4^, and these vocal responses are often accompanied by additional defensive behaviors^22–24^. Therefore, whether vocalization-related PAG neurons act as a discrete channel to selectively drive USV production or might also contribute to non-vocal motor behaviors remains unclear. We first confirmed that in addition to enhancing USV production, non-selective chemogenetic or optogenetic activation of caudolateral PAG neurons drove pronounced locomotor effects, including freezing and escape behaviors (Ext. Data Fig. 3, Movies S3-S4). In contrast, chemogenetic or optogenetic activation of CANE-tagged PAG-USV neurons elicited USVs without evoking any of these non-vocal behaviors (Ext. Data Fig. 3, Movies S5-6). These observations indicate that PAG-USV neurons are selectively involved in USV production and are intermingled with other neurons in the caudolateral PAG that contribute to various non-vocal behaviors^6–9^.

An influential idea is that neural activity in the PAG gates vocalization by initiating vocal output and specifying the duration of vocal bouts, rather that participating in the temporal patterning of individual vocal elements^2–4^. Here we tested this idea explicitly by comparing the temporal pattern of optogenetic stimulation of PAG-USV neurons to the temporal pattern of optogenetically-elicited USVs. In support of a vocal gating role for PAG-USV neurons, we found that the duration of a given vocal bout was on average similar to the duration of the laser stimulus used to elicit vocalization (Fig. 3D, N=6 PAG-ChR2 mice, N=2 PAG_USV_-ChR2 mice). We also observed that tonic (i.e., unpatterned) and 10 Hz laser stimuli both elicited bouts of multiple USV syllables (Fig. 3B,E) and that individual USVs elicited by tonic and 10Hz laser stimuli were of similar duration (Fig. 3F, 49.1 ± 27.3 s for tonic, 45.3 ± 19.9 s for 10 Hz, p=0.38, Wilcoxon signed-rank test). Thus, sustained PAG-USV activation is required for sustained vocal output, but patterned PAG-USV activation is not necessary for the production of bouts containing multiple USV syllables.

These findings do not exclude the possibility that patterned PAG-USV activity, when present, is capable of driving the timing of individual USV syllables. To evaluate this possibility, we examined whether the onsets of individual USVs elicited by 10 Hz laser stimuli were entrained to the timing of the immediately preceding laser pulse. Notably, USV onset times were not tightly entrained to a particular point in the laser duty cycle, indicating that the timing of PAG-USV activation does not precisely determine when individual USVs are initiated (Fig. 3G-H, n=645 USVs from PAG-ChR2 mice and n=306 USVs from PAGUSV-ChR2 mice). Instead, USV onsets were precisely aligned to the ongoing respiratory cycle, occurring just after the end of the inspiratory phase and during expiration, as has been demonstrated in freely vocalizing and moving rodents^25^ (Fig. 3G-H, n=648 opto-USVs measured in a head-fixed PAG_USV_-ChR2 mouse, see Methods). We also found subtle effects of PAG-USV activation on respiration, including an increase in the proportion of the respiratory cycle dedicated to expiration (Fig. 3I-J, p<0.001, Wilcoxon signed-rank test), consistent with observed changes in respiration that occur during USV production in freely behaving rodents^25^. Taken together, these findings support the idea that elevated PAG-USV neural activity is required to initiate and sustain a vocal bout, while the timing of individual USVs within the bout is controlled largely by neural circuits that govern the ongoing respiratory cycle.

A remaining issue is to determine the efferent targets of PAG-USV neurons, knowledge of which can ultimately shed light on how the PAG engages downstream circuits important to vocal patterning. Therefore, we used CANE to express GFP in PAG-USV neurons and mapped their axonal projections (N=4 mice, see Methods). We found that the axons of PAG-USV neurons terminate in the lateral parabrachial nucleus and in pontine and medullary reticular regions that have been speculated to contain premotor pattern-generating circuits important for vocal control^3^. These reticular targets of PAG-USV axons include the pontine reticular formation, the gigantocellular reticular formation of the rostral medulla, and the dorsal and ventral reticular formation of the caudal medulla, as well as other regions of the brainstem reticular formation (Fig. 4). Notably, we did not detect any GFP-labeled axons in nucleus ambiguus, which contains the laryngeal motor neurons, consistent with the idea that PAG-USV neurons are upstream of central pattern generating elements that directly engage vocal motor neurons. Indeed, axons of PAG-USV neurons terminate extensively in the nucleus retroambiguus (RAm), which contains premotor neurons that project to both laryngeal and expiratory motor neurons^3,12,26,27^ (Fig. 4G). Notably, USV production also is accompanied by high levels of Fos expression in RAm neurons, and this Fos expression was absent in muted mice in which PAG-USV neurons were silenced with TeLC (Ext. Data Fig. 4). Thus, PAG-USV neurons can excite downstream vocal-respiratory premotor neurons that are engaged during vocalization.

**Figure 4.**
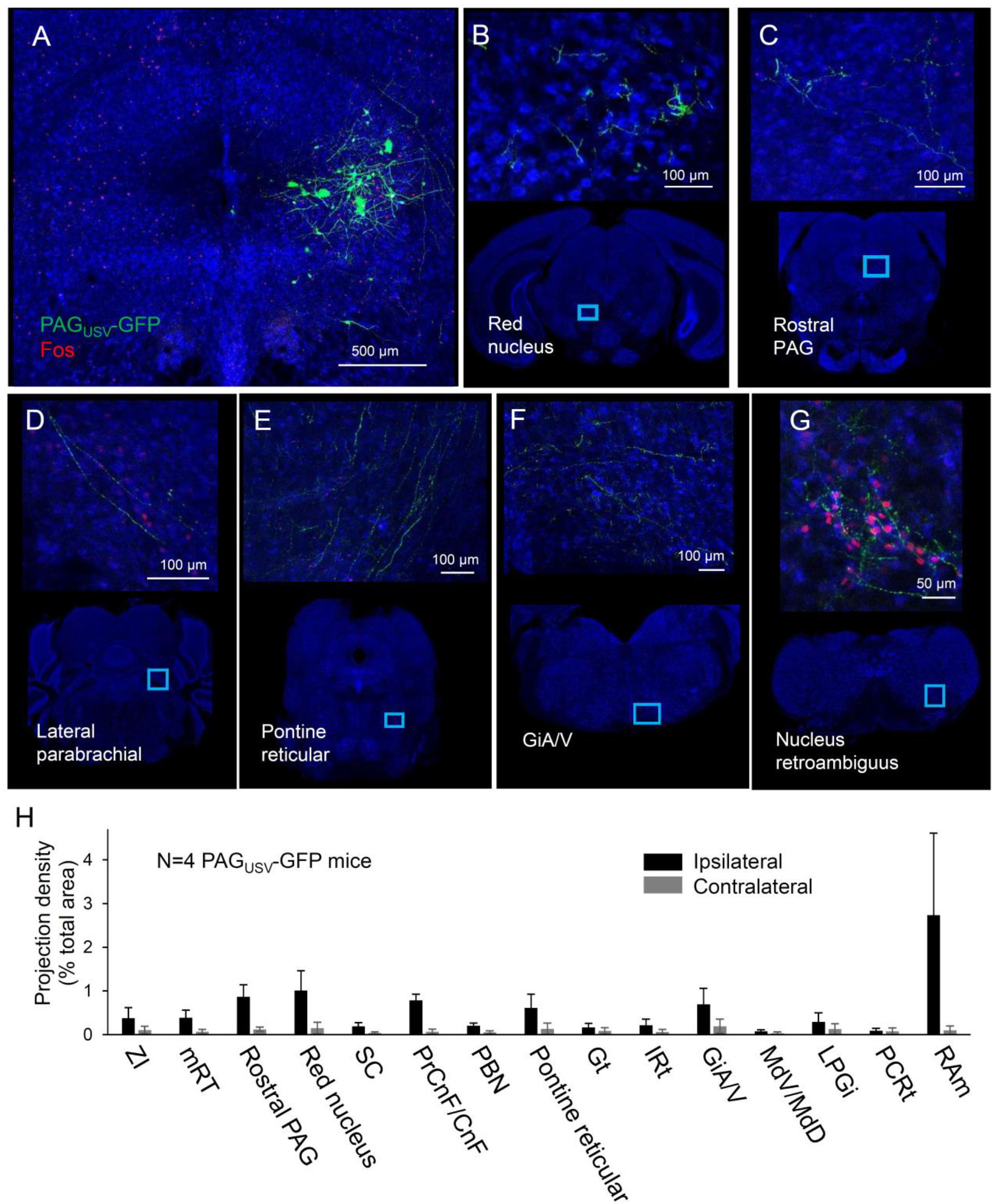
PAG-USV neurons project to brainstem vocal-motor and respiratory centers. Representative confocal images show (A) representative CANE-driven GFP labeling of PAG-USV cell bodies and PAG-USV axonal projections to (B) the red nucleus, (C) rostral PAG, (D) the lateral parabrachial nucleus, (E) the pontine reticular formation, (F) the magnocellular reticular formation of the rostral medulla, and (G) nucleus retroambiguus in the caudal medulla. (H) Quantification of axonal projections, presented as projection density (total square pixels of axonal innervation normalized by the total area of each region, see Methods, N=4 mice, mean ± SE). ZI, zona incerta; mRT, mesencephalic reticular formation; SC, superior colliculus; PrCnF, precuneiform; CnF, cuneiform; PBN, parabrachial nucleus; Gt, gigantocellular reticular formation; IRt, intermediate reticular formation; GiA/V, magnocellular reticular formation; MdV, ventral medullary reticular formation; MdD, dorsal medullary reticular formation; LPGi, paragigantocellular reticular formation; PCRt, parvocellular reticular formation; RAm, nucleus retroambiguus.

Here we used a novel viral genetic strategy to identify neurons in the midbrain PAG whose activity is essential for the production of courtship USVs in the male mouse. Notably, silencing PAG-USV neurons selectively abolishes USV production, while their artificial activation selectively drives vocalization in the absence of social cues, even though USVs are normally produced as part of a suite of coordinated courtship behaviors and only in response to female social cues^18,20,21^. Thus, the PAG contains a subset of neurons that are selectively involved in controlling USV output, adding to a growing body of evidence that the PAG contains anatomically distinct subsets of neurons specialized to drive specific behaviors important for reproduction and survival^8,28^. Moreover, by selectively suppressing the male’s ability to produce USVs while leaving his other courtship behaviors intact, we established the salience of male USVs for female social preference and affiliation. Selective optogenetic activation of PAG-USV neurons revealed that their sustained activation is required for sustained vocal output, but that the pattern of elicited USVs is entrained to ongoing respiration and not to the pattern of PAG-USV activation. These findings support the idea that PAG-USV neurons integrate sensory and motivational inputs to subsequently trigger downstream vocal premotor neurons to drive socially-appropriate USV production. In direct support of this idea, we found that PAG-USV axons terminate in brainstem regions that contain vocal and respiratory premotor neurons and that are thought to contain vocal pattern generators. Although how and where vocal commands from the PAG are integrated with the ongoing respiratory rhythm remains unknown, an intriguing possibility is that input from respiratory brainstem regions^29,30^ could rhythmically gate the responsiveness of vocal premotor neurons to PAG-USV input, such that the PAG can only excite vocal motor neuron pools during expiration. Taken together, our findings establish that PAG-USV neurons provide an essential and selective pathway for vocal control, affording an entry point for exploring the brain-wide circuits for vocal communication.

## Methods

### Animal statement

All experiments were conducted according to protocols approved by the Duke University Institutional Animal Care and Use Committee.

### Animals

Adult (P60-90) male Fos^TVA^ mice (Jackson Laboratory, stock 027831) were used for capturing PAG-USV neurons with the CANE method^10,11^. Fos^TVA^ mice were housed on a reversed light/dark cycle and were single-housed beginning ~7 days prior to experimentation. Vglut2-ires-Cre (Jackson Laboratory, stock 028863) and PV-Cre (Jackson Laboratory stock 008069) group-housed adult males were used to express hM3Dq and ChR2 respectively in caudolateral PAG neurons.

### USV recording and analysis

To elicit USVs for subsequent Fos immunohistochemistry, single-housed males were presented with either a freely moving female, a female restrained in a small container that still permitted limited olfactory investigation by the male, or female urine (collected and pooled from multiple females immediately before test session, presented 2-4 times during a 60 min. session). USVs were recorded with an ultrasonic microphone (Avisoft, CMPA/CM16), amplified (Presonus TubePreV2), and digitized at 250 kHz (Spike 7, CED). USVs were detected using codes from the Holy lab (http://holylab.wustl.edu/) using the following parameters (mean frequency > 45 kHz; spectral purity > 0.3; spectral discontinuity < 0.85; min. USV duration = 5 ms; minimum inter-syllable interval = 30 ms). To elicit USVs for tagging of PAG-USV neurons using CANE, Fos^TVA^ males were given social experience with a female (30–60 min. session), either in their home cage fitted with an acoustically permeable lid or in a test chamber that had no lid and allowed easy microphone access. Sixty minutes from the start of the session, Fos^TVA^ males were anesthetized and taken for injection of the PAG with viruses (see below), such that injections began approximately 2 hours from the start of USV production.

### Fos immunohistochemistry

Mice were deeply anaesthetized with isoflurane and then transcardially perfused with ice-cold 4% paraformaldehyde in 0.1 M phosphate buffer, pH 7.4 (4% PFA). Dissected brain samples were then post-fixed overnight in 4% PFA at 4 °C, cryoprotected in a 30% sucrose solution in PBS at 4 °C for 48 hours, frozen in Tissue-Tek O.C.T. Compound (Sakura), and stored at –80 °C until sectioning. Brains were cut into 80 μm coronal sections, rinsed 3x in PBS, permeabilized for 3 hours in PBS containing 1% Triton X (PBST), and then blocked in 0.3% PBST containing 10% Blocking One (Nacalai Tesque; i.e., blocking solution). Sections were then processed for 48 hours at 4 degrees with the primary antibody in blocking solution (1:400 goat anti-Fos, Santa Cruz, sc52-g), rinsed 3 × 10 mins. in PBS, then processed for 48 hours at 4 degrees with secondary antibodies in blocking solution (1:1000, Alexa Fluor 488 or 594 bovine anti-goat, Jackson Laboratories, plus 1:500 NeuroTrace, Invitrogen). Tissue sections rinsed again 3 × 10 mins. in PBS, mounted on slides, and coverslipped with Fluoromount-G (Southern Biotech). After drying, slides were imaged with a 10x objective on a Zeiss 700 laser scanning confocal microscope, and Fos-positive neurons were counted manually.

### Viruses and expression of transgenes in PAG-USV neurons using CANE method

To stably express transgenes in PAG-USV neurons, Fos^TVA^ males were given social experience with a female (30-60 mins.) that resulted in high levels of USV production (500-5000 USVs total). Males were then anesthetized (1.5-2% isoflurane), and the caudolateral PAG was targeted for injection. The final injection coordinates were AP=-4.7 +/-0.2 mm, ML= 0.7, DV= 1.75, and these were reached by making a small craniotomy at approximately AP=-3.3 +/-0.2 mm, ML=0.7 and advancing the pipette back toward PAG at a 30 degree angle relative to vertical. The PAG was then injected with a mixture of CANE-lenti-Cre and an AAV driving Cre-dependent expression of the transgene of interest (1:1 ratio, 400-500 nL total, pressure-injected with a Nanoject II (Drummond), 4.6 nL every 15s). CANE-lenti-Cre was produced as previously described^10,11^. The following AAVs were co-injected with CANE-lenti-Cre: AAV2/1-CAG-FLEX-GFP (UNC Vector Core), AAV2/8-hSyn-FLEX-GFP (UNC Vector Core), AAV2/8-hSyn-FLEX-TeLC-P2A-GFP (produced as previously described^31^), AAV-hSyn-FLEX-hM3Dq-mCherry (Addgene and also UNC Vector Core), and AAV2/1-CBA-FLEX-ChR2-mCherry (UPenn Vector Core). To non-selectively drive expression of ChR2 in caudolateral PAG neurons, PV-Cre males were injected unilaterally in the caudolateral PAG with AAV2/1-hSyn-ChR2-eYFP (UPenn Vector Core, 50 nL). To drive expression of hM3Dq in caudolateral PAG neurons, VGlut2-Cre males were injected unilaterally in the caudolateral PAG with AAV2/8-hSyn-FLEX-hM3Dq-mCherry (50 nL). The waiting times following virus injections were as follows: CANE-lenti-Cre plus AAV-FLEX-GFP, 4 weeks; CANE-lenti-Cre plus AAV-FLEX-TeLC, 2 weeks; CANE-lenti-Cre plus AAV-FLEX-hM3Dq, 6 weeks; CANE-lenti-Cre plus AAV-FLEX-ChR2, 4 weeks; AAV-ChR2 only, 3 weeks; AAV-FLEX-hM3Dq only, 6 weeks.

### Quantification of courtship behavior

Time spent engaged in courtship behavior by either PAG_USV_-TeLC males or control males was manually scored from videos recorded with a webcam (Logitech, 30 frames per second) in which the male was allowed to freely interact with a novel female placed in his home cage (30 min. session). Anogenital sniffing and following were scored as low intensity courtship behaviors, while mounting was scored as a high intensity courtship behavior.

### Three-chambered social preference test

During an acclimation period, the female test mouse was placed in the central chamber and allowed to freely explore the three chambers for approximately 5 minutes. The female was then restricted within the central chamber, and a male was placed in each side chamber, restrained underneath a wire cage insert that permitted limited olfactory investigation of the female when she was nearby. The female was then allowed to move freely between the three chambers, and her preference for the PAG_USV_-GFP control male was measured as (time spent on side with control male)/(time spent with control male + time spent with muted PAGUSV-TeLC male). An ultrasonic microphone was placed above each of the two side chambers, and we determined empirically that the vast majority of USVs produced by the control PAG_USV_-GFP could not be detected or appeared at much lower amplitude on the microphone on the opposite side of the chamber, thus allowing us to unambiguously assign all USVs as arising from the control male’s side of the chamber. Although we cannot completely rule out the possibility that the female test mouse contributed to the total USV count, it has been reported that female mice vocalize to males when they are being actively courted and when the male is also vocalizing^21^. Even under these circumstances, females were reported to contribute ~20% of the total measured USVs^21^, and we never observed overlapping USVs that would indicate that the control male and the female were vocalizing at the same time. Female mice were of a variety of genotypes, always 2-3 months of age (to ensure that high frequency hearing was intact), and estrous state was not measured or controlled.

### CNO dosage and delivery

PAG_USV_-hM3Dq mice were lightly anesthetized with isoflurane and then injected i.p. with CNO (4 mg/kg from Sigma, or 15 mg/kg from Duke Small Molecule Synthesis Facility) or an equivalent volume of sterile saline. PAG-hM3Dq mice were injected with 1 mg/kg CNO (Duke Small Molecule Synthesis Facility) or an equivalent volume of sterile saline. CNO-only control mice were anesthetized lightly and injected i.p. with either 4 mg/kg CNO (Sigma) or 15 mg/kg CNO (Duke Small Molecule Synthesis Facility).

### Optogenetic stimulation of PAG neurons

Custom-made optogenetic ferrules were implanted bilaterally (PAG_USV_-ChR2 mice, implanted 3-4 weeks following viral injection) or unilaterally (PAG-ChR2 mice, implanted in the same surgery as viral injection) above the caudolateral PAG at least 1 week before the first optogenetic testing and were fixed to the skull using Metabond (Parkell). PAG neurons were optogenetically activated with illumination from a 473 nm laser (3-30 mW) at either 10 Hz (50 ms pulses, 1-2s total) or with phasic laser pulses (0.05-2s duration). Laser stimuli were driven by computer-controlled voltage pulses (Spike 7, CED).

### Quantification of optogenetically-elicited body movements

The mouse’s position was measured using custom Matlab codes (K. Tschida) that detected and tracked the centroid of the mouse’s body position across video frames (Logitech webcam, 30 frames per second), and speed of movement was calculated as the change in position across pairs of frames. To align movement with optogenetic activation of PAG neurons, we first estimated the temporal offset between the webcam video and USV audio by calculating the time of the peak cross-covariance between the high-pass filtered webcam audio and the low-pass filtered USV audio. This offset was then used to align the mouse’s movement to the onset of each optogenetic laser stimulus.

### Comparison of acoustic features of chemogenetically- and optogenetically-elicited USVs to female-directed USVs

We used custom Matlab codes to generate spectrograms of our audio recordings and manually annotated onset and offset times of all opto-USVs. For all annotated USVs, we then calculated 5 acoustic features: (1) duration, (2) inter-syllable interval (defined as interval from start of one USV to start of the next USV, >1000 ms intervals not included in analysis), (3) mean pitch (dominant frequency calculated at each time point of the USV, then averaged across entire syllable), (4) pitch variance (defined as the variance of the dominant frequency for a syllable), and (5) amplitude (defined as bandpower from 30-125 kHz, converted to dB, and measured relative to quiet background noise in recording). These measurements of acoustic features were compared to those taken from female-directed syllables produced by the same Fos^TVA^ males, as well as 15 control Fos^TVA^ males. Female-directed USVs were automatically detected using Matlab codes (http://holylab.wustl.edu/), but we also manually annotated and analyzed the acoustic features of a subset of female-directed USVs to ensure that any observed differences between conditions could not be attributed to the detection method.

### Respiration measurements and analyses

An air flow sensor (Honeywell, AMW3300V) was placed as close as possible to the snout of a head-fixed PAGUSV-ChR2 male that was allowed to run on a treadmill. Custom Matlab codes (K. Tschida) were used to detect inspiration onset and expiration onset throughout the respiratory cycle. Briefly, breathing data were collected at 250 kHz (Spike 7, CED), downsampled to 1 kHz, and the portion of the breathing signal to be analyzed was detrended, mean-subtracted, and divided by its own standard deviation. The breathing signal was then inverted and converted into a binary vector (values>0 set to 1 and values<0 set to 0). Inspiration onset times were detected as signal crossings from 0 to 1, and inspiration offsets were detected as signal crossings from 1 to 0. Respiration rate over a given interval was calculated in Hz as 1/(mean time between inspiration onsets).

### GFP labeling of PAG-USV neurons and axon projection tracing

PAG-USV neurons were labeled with GFP by injecting the caudolateral PAG of a Fos^TVA^ male with a mixture of CANE-lenti-Cre and AAV-FLEX-GFP following a vocal encounter with a female. After at least 4 weeks, PAG_USV_-GFP males were sacrificed, their brains were processed as detailed above, and every other 80 micron-thick tissue section was imaged with a 10x objective on a Zeiss 700 laser scanning confocal microscope. To detect axons in each image, images were binarized using a custom Matlab code (K. Tschida, adapted from code provided by J. Takatoh). Brain regions were then manually annotated (ImageJ), and the area in square pixels containing axons was calculated for each brain region for each mouse and then normalized by the area of that brain region. Brain regions with an innervation density of less than 0.07% or that were not observed to be innervated in 4/4 mice were excluded from analysis.

### Data availability

The data that support the findings of this study are available from the corresponding author upon reasonable request.

### Code availability

All custom-written Matlab codes used in this study are available from the corresponding author.

### Statistics

Non-parametric, two-sided statistical comparisons were used in all analyses (Mann Whitney to compare two groups of unpaired observations, Wilcoxon signed-rank for paired observations, alpha=0.05), with the exception of the comparison of time spent immobile for PAG-hM3Dq and PAG_USV_-hM3Dq mice treated with saline and CNO (Ext. Data Fig. 3A), for which we used a two-way repeated measures ANOVA followed by post-hoc paired t-tests. No statistical methods were used to predetermine sample sizes. Error bars represent standard error of the mean unless otherwise noted.

## Supplementary Information

We have included six supplemental movies, with legends included after the main and extended data figures and legends.

**Extended Data Figure 1.**
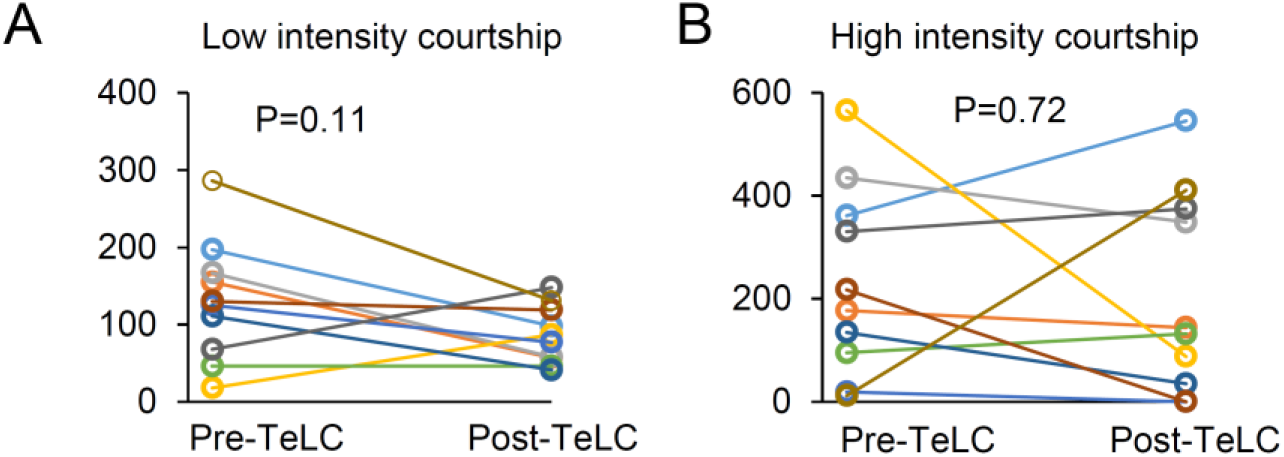
Blocking transmitter release from PAG-USV neurons has no effect on the amount of time males engage in high intensity or low intensity courtship behaviors. (A) The amount of time males spend in low intensity courtship (anogenital sniffing and following) is not changed following silencing of PAG-USV neurons (N=10 males, p=0.11, Wilcoxon ranked-sign test). (B) Same, for high intensity courtship (i.e., mounting) (p=0.72).

**Extended Data Figure 2.**
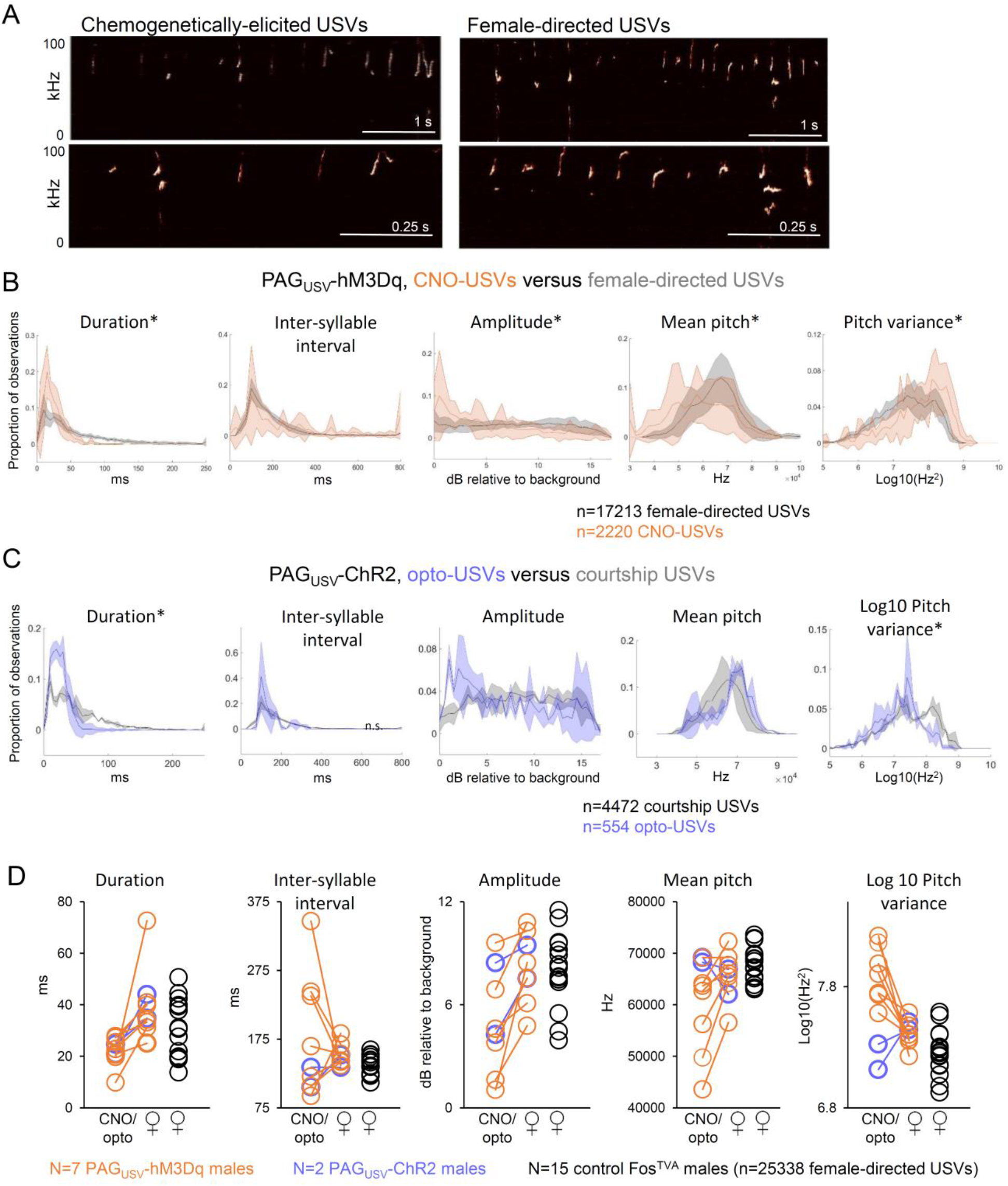
Comparison of acoustic features of chemogenetically- and optogenetically-elicited USVs versus courtship USVs. (A) Spectrograms show CNO-USVs (left) and female-directed USVs (right) from the same PAG_USV_-hM3Dq male. Bottom panels show expanded view from top panels. (B) Distributions of 5 acoustic parameters are shown for CNO-USVs (orange) and female-directed USVs (gray) for 7 PAG_USV_-hM3Dq mice (lines show mean values, shading represents standard deviation). Asterisks indicate acoustic parameters whose median value changed significantly in the same direction for at least 6 of 7 mice (defined as p<0.05 for Mann Whitney U tests performed for a given mouse and acoustic parameter). (C) Distributions of 5 acoustic parameters are shown for opto-USVs (blue) and courtship USVs (gray) for 2 PAG_USV_-ChR2 mice. Asterisks indicate acoustic parameters whose median value changed significantly in the same direction for both mice (p<0.05, Mann Whitney U tests.) (D) Median values of 5 acoustic parameters for CNO/opto-USVs and courtship USVs from PAG_USV_-hM3Dq mice (orange connected points) and PAG_USV_-ChR2 mice (blue connected points) are compared to median values measured from 15 control Fos^TVA^ mice producing courtship USVs (black).

**Extended Data Figure 3.**
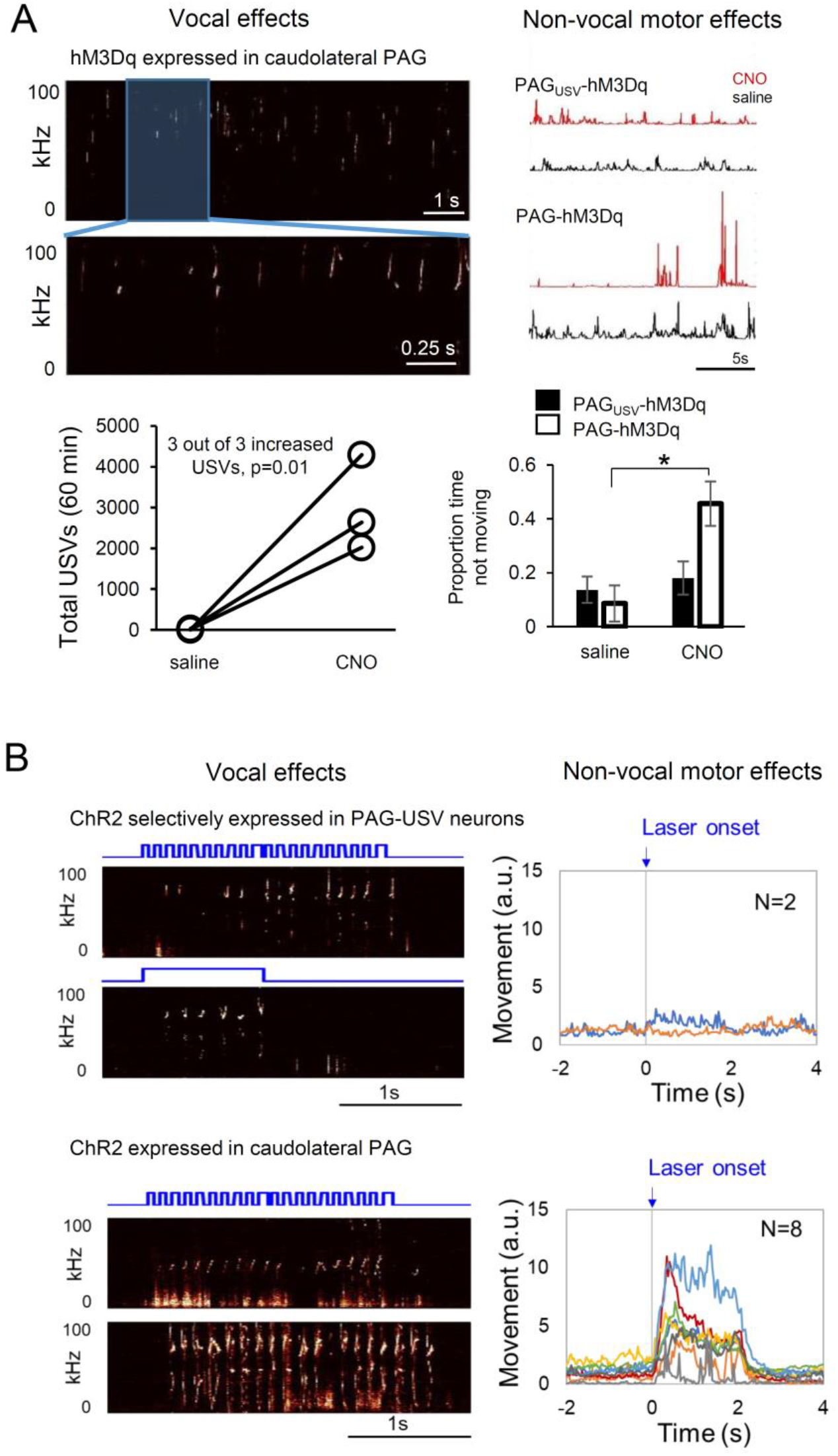
Non-selective chemogenetic and optogenetic activation of caudolateral PAG neurons drives USV production as well as non-vocal movements. (A) (Top, left) Spectrograms show USVs produced by a VGlut-Cre male after injection of AAV-FLEX-hM3Dq bilaterally into the caudolateral PAG and treatment with CNO. (Bottom, left) All 3 VGlut-Cre males with hM3Dq expression in the caudolateral PAG dramatically increased USV production following treatment with CNO as compared to saline. (Top, right) Representative movement traces from a PAG_USV_-hM3Dq male and a PAG-hM3Dq male, after treatment with either CNO (red traces) or saline (black traces). Note that the movement of the PAG-hM3Dq male after CNO treatment consists of periods of immobility punctuated by periods of high speed movement. (Bottom, right) Quantification of proportion of time spent immobile (>1s periods) for PAGusv-hM3Dq (N=8, black bars) and PAG-hM3Dq males (N=3, white bars) after treatment with either saline or CNO (two-way repeated measures ANOVA revealed significant interaction between group and treatment, p=0.01 for PAG-hM3Dq CNO versus saline, p=0.28 for PAG_USV_-hM3Dq CNO versus saline, paired t-tests, mean ± SE are shown). (B) (Top, left) Selective expression of ChR2 in PAG-USV neurons drives USV production during laser stimulation (top, left) but not non-vocal movements (top, right). In contrast, non-selective ChR2 expression of ChR2 in caudolateral PAG neurons drives both USV production (bottom, left) and high speed movements (bottom, right) during laser stimulation (N=8 mice). Please note that spectrograms from PAGUSV-ChR2 mice are reproduced from those in Fig. 3B.

**Extended Data Figure 4.**
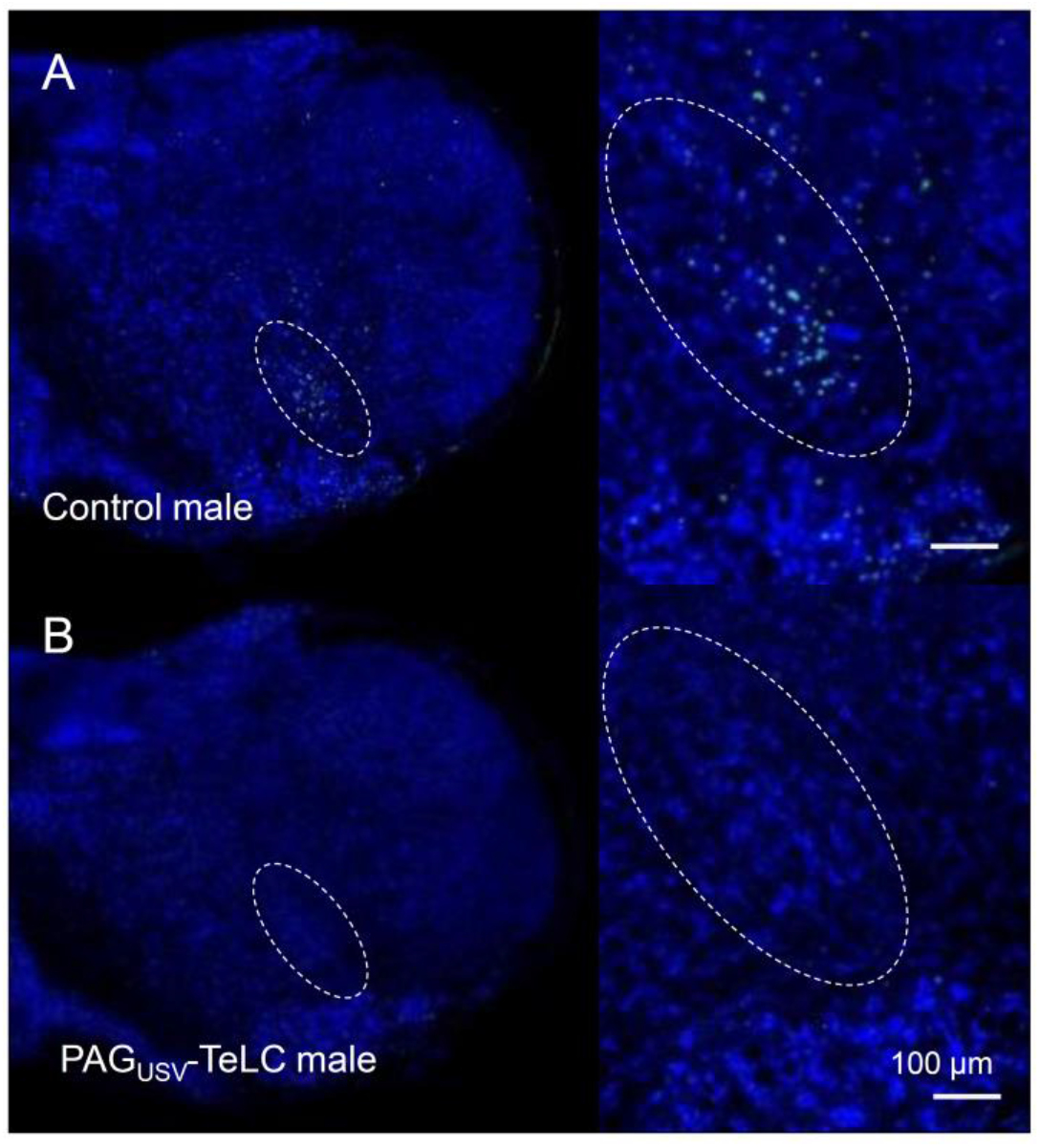
Silencing of PAG-USV neurons blocks USV-induced Fos expression in the nucleus retroambiguus. (A) USV-induced Fos expression (green) in RAm is shown for a control male (inset shows magnified view; blue, NeuroTrace). USV-induced Fos in RAm was observed in 10/11 control males. (B) USV-induced Fos expression is absent in a male expressing TeLC in his PAG-USV neurons. USV-induced Fos in RAm was observed in 0/5 PAG_USV_-TeLC males.

**Movie S1. Male mouse courting a female and producing courtship USVs.** A Fos^TVA^ male is shown courting a female prior to injection of viruses to drive TeLC expression in PAG-USV neurons. Video is shown at the top, a spectrogram (bottom) showing the male’s courtship USVs is synchronized to the video, and pitch-shifted audio (80 kHz to 5 kHz transformation) is included to place the courtship USVs within the human hearing range.

**Movie S2. Blocking neurotransmitter release in PAG-USV neurons abolishes the production of male courtship USVs without affecting non-vocal courtship.** Same as Movie S1, except the same Fos^TVA^ male is now shown courting a female following bilateral expression of TeLC in his PAG-USV neurons (~14 days post-injection). Note that although he still exhibits typical non-vocal courtship behaviors, he is unable to produce courtship USVs.

**Movies S3: Non-selective chemogenetic activation of caudolateral PAG neurons promotes USV production, in addition to scurrying and immobility.** Treatment with CNO (i.p.) promotes USV production, as well as non-vocal movements punctuated by periods of immobility, in a male mouse expressing hM3Dq in caudolateral PAG neurons.

**Movie S4: Non-selective optogenetic activation of the caudolateral PAG neurons drives USV production, in addition to robust locomotor effects:** Blue laser light (10 Hz, 473 nm) optogenetically activates ChR2-expressing caudolateral PAG neurons and elicits both USV production and non-vocal movements

**Movie S5. Selective chemogenetic activation of PAG-USV neurons increases USV production in the absence of any social cues.** A Fos^TVA^ male with bilateral CANE-driven expression of hM3Dq in PAG-USV neurons produces USVs after treatment with CNO (i.p.).

**Movie S6: Selective optogenetic activation of PAG-USV neurons drives USV production in the absence of any social cues:** Blue laser light (10 Hz, 473 nm) is used to optogenetically activate ChR2-expressing PAG-USV neurons and elicit USVs in a Fos^TVA^ male, tested alone in the absence of any social partner or female cues.

## Acknowledgements

Thanks to Jun Takatoh for providing Matlab code to detect GFP-positive axons, and to Michael Booze for additional mouse husbandry.

## Author contributions

RM, FW, and KAT designed the experiments. KAT and VM conducted the experiments. SZ made viruses for the experiments. KS provided guidance in the use of the CANE method and made viruses used in the experiments. BXH provided animal husbandry and experimental support. RM, FW, and KAT wrote the manuscript, and all authors approved the final manuscript.

## Data deposition statement

The data that support the findings of this study are available from the corresponding author upon reasonable request. All custom-written Matlab codes used in this study are available from the corresponding author.

## References

1 Guenther, F. H. Neural control of speech.

2 Jurgens, U. The neural control of vocalization in mammals: a review. J Voice 23, 1–10, doi:10.1016/j.jvoice.2007.07.005 (2009).

3 Jurgens, U. Neural pathways underlying vocal control. Neurosci Biobehav Rev 26, 235–258 (2002).

4 Jurgens, U. The role of the periaqueductal grey in vocal behaviour. Behav Brain Res 62, 107–117 (1994).

5 Esposito, A., Demeurisse, G., Alberti, B. & Fabbro, F. Complete mutism after midbrain periaqueductal gray lesion. Neuroreport 10, 681–685 (1999).

6 Bandler, R. & Shipley, M. T. Columnar organization in the midbrain periaqueductal gray: modules for emotional expression? Trends Neurosci 17, 379–389 (1994).

7 Carrive, P. The periaqueductal gray and defensive behavior: functional representation and neuronal organization. Behav Brain Res 58, 27–47 (1993).

8 Tovote, P. et al. Midbrain circuits for defensive behaviour. Nature 534, 206–212, doi:10.1038/nature17996 (2016).

9 Holstege, G. The periaqueductal gray controls brainstem emotional motor systems including respiration. Prog Brain Res 209, 379–405, doi:10.1016/B978-0-444-63274-6.00020-5 (2014).

10 Sakurai, K. et al. Capturing and manipulating activated neuronal ensembles with CANE delineates a hypothalamic social fear circuit. Neuron In press (2016).

11 Rodriguez, E. et al. A craniofacial-specific monosynaptic circuit enables heightened affective pain. Nat Neurosci 20, 1734–1743, doi:10.1038/s41593-017-0012-1 (2017).

12 Holstege, G. Anatomical study of the final common pathway for vocalization in the cat. J Comp Neurol 284, 242–252, doi:10.1002/cne.902840208 (1989).

13 Subramanian, H. H. & Holstege, G. The nucleus retroambiguus control of respiration. J Neurosci 29, 3824–3832, doi:10.1523/JNEUROSCI.0607-09.2009 (2009).

14 Holy, T. E. & Guo, Z. Ultrasonic songs of male mice. PLoS Biol 3, e386, doi:10.1371/journal.pbio.0030386 (2005).

15 Whitney, G., Alpern, M., Dizinno, G. & Horowitz, G. Female odors evoke ultrasounds from male mice. Animal Learning and Behavior 2, 13–18 (1974).

16 Nyby, J., Wysocki, C. J., Whitney, G., Dizinno, G. & Schneider, J. Elicitation of male mouse (Mus muscululs) ultrasonic vocalization: 1. Urinary Cues. Journal of Comparative and Physiological Psychology 93, 957–975 (1979).

17 Gourbal, B. E., Barthelemy, M., Petit, G. & Gabrion, C. Spectrographic analysis of the ultrasonic vocalisations of adult male and female BALB/c mice. Naturwissenschaften 91, 381–385, doi:10.1007/s00114-004-0543-7 (2004).

18 Nyby, J. Ultrasonic vocalizations during sex behavior of male house mice (Mus musculus): a description. Behav Neural Biol 39, 128–134 (1983).

19 White, N. R., Prasad, M., Barfield, R. J. & Nyby, J. G. 40- and 70-kHz vocalizations of mice (Mus musculus) during copulation. Physiol Behav 63, 467–473 (1998).

20 Matsumoto, Y. K. & Okanoya, K. Phase-Specific Vocalizations of Male Mice at the Initial Encounter during the Courtship Sequence. PLoS One 11, e0147102, doi:10.1371/journal.pone.0147102 (2016).

21 Neunuebel, J. P., Taylor, A. L., Arthur, B. J. & Egnor, S. E. Female mice ultrasonically interact with males during courtship displays. Elife 4, doi:10.7554/eLife.06203 (2015).

22 Waldbillig, R. J. Attack, eating, drinking, and gnawing elicited by electrical stimulation of rat mesencephalon and pons. J Comp Physiol Psychol 89, 200–212 (1975).

23 Magoun, H. W., Atlas, D., Ingersoll, E. H. & Ranson, S. W. Associated Facial, Vocal and Respiratory Components of Emotional Expression: An Experimental Study. J Neurol Psychopathol 17, 241–255 (1937).

24 Bandler, R. & Carrive, P. Integrated defence reaction elicited by excitatory amino acid microinjection in the midbrain periaqueductal grey region of the unrestrained cat. Brain Res 439, 95–106 (1988).

25 Sirotin, Y. B., Costa, M. E. & Laplagne, D. A. Rodent ultrasonic vocalizations are bound to active sniffing behavior. Front Behav Neurosci 8, 399, doi:10.3389/fnbeh.2014.00399 (2014).

26 VanderHorst, V. G., Terasawa, E. & Ralston, H. J., 3rd. Monosynaptic projections from the nucleus retroambiguus region to laryngeal motoneurons in the rhesus monkey. Neuroscience 107, 117–125 (2001).

27 Vanderhorst, V. G., Terasawa, E., Ralston, H. J., 3rd & Holstege, G. Monosynaptic projections from the nucleus retroambiguus to motoneurons supplying the abdominal wall, axial, hindlimb, and pelvic floor muscles in the female rhesus monkey. J Comp Neurol 424, 233–250 (2000).

28 Caggiano, V. et al. Midbrain circuits that set locomotor speed and gait selection. Nature 553, 455–460, doi:10.1038/nature25448 (2018).

29 Gerrits, P. O. & Holstege, G. Pontine and medullary projections to the nucleus retroambiguus: a wheat germ agglutinin-horseradish peroxidase and autoradiographic tracing study in the cat. J Comp Neurol 373, 173–185, doi:10.1002/(SICI)1096-9861(19960916)373:2<173::AID-CNE2>3.0.œ;2-0 (1996).

30 Song, G., Wang, H., Xu, H. & Poon, C. S. Kolliker-Fuse neurons send collateral projections to multiple hypoxia-activated and nonactivated structures in rat brainstem and spinal cord. Brain Struct Funct 217, 835–858, doi:10.1007/s00429-012-0384-7 (2012).

31 Zhang, Y. et al. Identifying local and descending inputs for primary sensory neurons. J Clin Invest 125, 3782–3794, doi:10.1172/JCI81156 (2015).

